# Population Bottlenecks and Intra-host Evolution during Human-to-Human Transmission of SARS-CoV-2

**DOI:** 10.1101/2020.06.26.173203

**Authors:** Daxi Wang, Yanqun Wang, Wanying Sun, Lu Zhang, Jingkai Ji, Zhaoyong Zhang, Xinyi Cheng, Yimin Li, Fei Xiao, Airu Zhu, Bei Zhong, Shicong Ruan, Jiandong Li, Peidi Ren, Zhihua Ou, Minfeng Xiao, Min Li, Ziqing Deng, Huanzi Zhong, Fuqiang Li, Wen-jing Wang, Yongwei Zhang, Weijun Chen, Shida Zhu, Xun Xu, Xin Jin, Jingxian Zhao, Nanshan Zhong, Wenwei Zhang, Jincun Zhao, Junhua Li, Yonghao Xu

## Abstract

The emergence of the novel human coronavirus, SARS-CoV-2, causes a global COVID-19 (coronavirus disease 2019) pandemic. Here, we have characterized and compared viral populations of SARS-CoV-2 among COVID-19 patients within and across households. Our work showed an active viral replication activity in the human respiratory tract and the co-existence of genetically distinct viruses within the same host. The inter-host comparison among viral populations further revealed a narrow transmission bottleneck between patients from the same households, suggesting a dominated role of stochastic dynamics in both inter-host and intra-host evolutions.

**Author summary:** In this study, we compared SARS-CoV-2 populations of 13 Chinese COVID-19 patients. Those viral populations contained a considerable proportion of viral sub-genomic messenger RNAs (sgmRNA), reflecting an active viral replication activity in the respiratory tract tissues. The comparison of 66 identified intra-host variants further showed a low viral genetic distance between intra-household patients and a narrow transmission bottleneck size. Despite the co-existence of genetically distinct viruses within the same host, most intra-host minor variants were not shared between transmission pairs, suggesting a dominated role of stochastic dynamics in both inter-host and intra-host evolutions. Furthermore, the narrow bottleneck and active viral activity in the respiratory tract show that the passage of a small number of virions can cause infection. Our data have therefore delivered a key genomic resource for the SARS-CoV-2 transmission research and enhanced our understanding of the evolutionary dynamics of SARS-CoV-2.

## Introduction

The rapid spread of the novel human coronavirus, SARS-CoV-2, has been causing millions of COVID-19 (coronavirus disease 2019) cases with high mortality rate worldwide [1,2]. As an RNA virus, SARS-CoV-2 mutates frequently due to the lack of sufficient mismatch repairing mechanisms during genome oreplication [3], leading to the development of genetically different viruses within the same host. Several studies have reported intra-host single nucleotide variants (iSNVs) in SARS-CoV-2 [4–6]. Recently, we investigated the intra-host evolution of SARS-CoV-2 and revealed genetic differentiation among tissue-specific populations [7]. However, it is still not clear how the intra-host variants circulate among individuals. Here, we described and compared viral populations of SARS-CoV-2 among COVID-19 patients within and across households. Our work here demonstrated the utilization of viral genomic information to identify transmission linkage of this virus.

## Results and discussion

Using both metatranscriptomic and hybrid-capture based techniques, we newly deep sequenced (respiratory tract (RT) samples of seven COVID-19 patients in Guangdong, China, including two pairs of patients from the same households, respectively (P03 and P11; P23 and P24). The data were then combined with those of 23 RT samples used in our previous study [7], yielding a combined data set of 30 RT samples from 13 COVID-19 patients (**Table S1**).

A sustained viral population should be supported by an active viral replication [8]. We firstly estimated the viral transcription activity within RT samples using viral sub-genomic messenger RNAs (**sgmRNAs**), which is only synthesised in infected host cells [9]. The sgmRNA abundance was measured as the ratio of short reads spanning the transcription regulatory sequence (TRS) sites to the viral genomic reads. The sgmRNA abundance within nasal and throat swab samples was similar to that within sputum samples (**Figure 1a**), reflecting an active viral replication in the upper respiratory (tract. Notably, the patient P01, who eventually passed away due to COVID-19, showed the highest level of sgmRNA abundance (**Figure S1**). Among the samples from patients with improved clinical outcomes, their viral Ct (cycle threshold) value of reverse transcriptase quantitative PCR (RT-qPCR) (negatively correlated with the days post symptoms onset (**Figure 1b**). Interestingly, the sgmRNA abundance showed a similar trend across time (**Figure 1c**). This result is further strengthened by the positive correlation between sgmRNA abundance and the Ct value (**Figure 1d**), reflecting a direct biological association between viral replication and viral shedding in the respiratory tract tissues.

**Figure 1.**
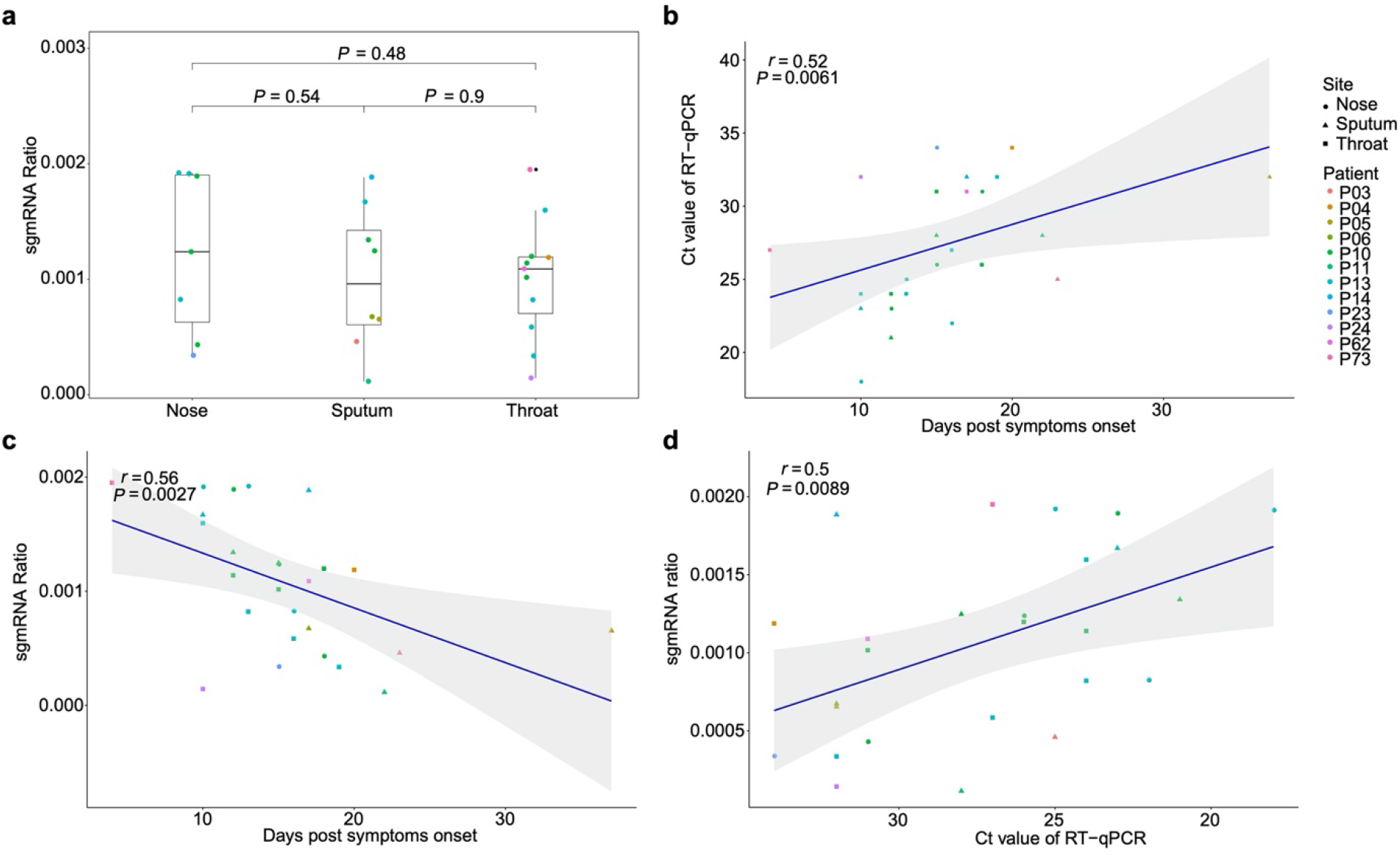
sub-genomic messenger RNAs (sgmRNAs) **a,** The ratio of sgmRNA of each respiratory sample type (nasal, throat swabs and sputum)**. b,** Correlation between the cycle threshold (Ct) of RT-qPCR and the days post symptoms onset. **c,** Correlation between estimated sgmRNA ratio and the days post symtoms onset. **d,** Correlation between estimated sgmRNA ratio and the cycle threshold of RT-qPCR.

Using the metatranscriptomic data, we identified 66 iSNVs in protein encoding regions with the (alternative allele frequency (AAF) ranged from 5% to 95% (**Table S2 and Table S3**). The identified iSNVs showed a high concordance between the AAFs derived from metatranscriptomic and that from (hybrid-capture sequences (Spearman’s *ρ* = 0.81, *P* < 2.2e-16; **Figure S2**). We firstly looked for signals of natural selection against intra-host variants. Using the Fisher’s exact test, we compared the number of iSNV sites on each codon position against that of the other two positions and detected a (significant difference among them (codon position 1[n = 10, *P* = 0.02], 2 [n = 21; *P* = 1] and 3 [n = 35; *P* = 0.03]). However, those iSNVs did not show a discriminated AAF among the non-synonymous and synonymous categories (**Figure 2a**), suggesting that most non-synonymous variants were not under an effective purifying selection within the host. Among the 66 identified iSNVs, 30 were coincided with the consensus variants in the public database (**Table S2**). Those iSNVs were categorised into common iSNVs, while the iSNVs presented in a single patient were categorised into rare iSNVs. Interestingly, the common iSNVs had a significant higher minor allele frequency compared to the rare iSNVs (**Figure S3;** Wilcoxon rank sum test, *P* = 2.7e-05), suggesting that they may have been developed in earlier strains before the most recent infection.

**Figure 2.**
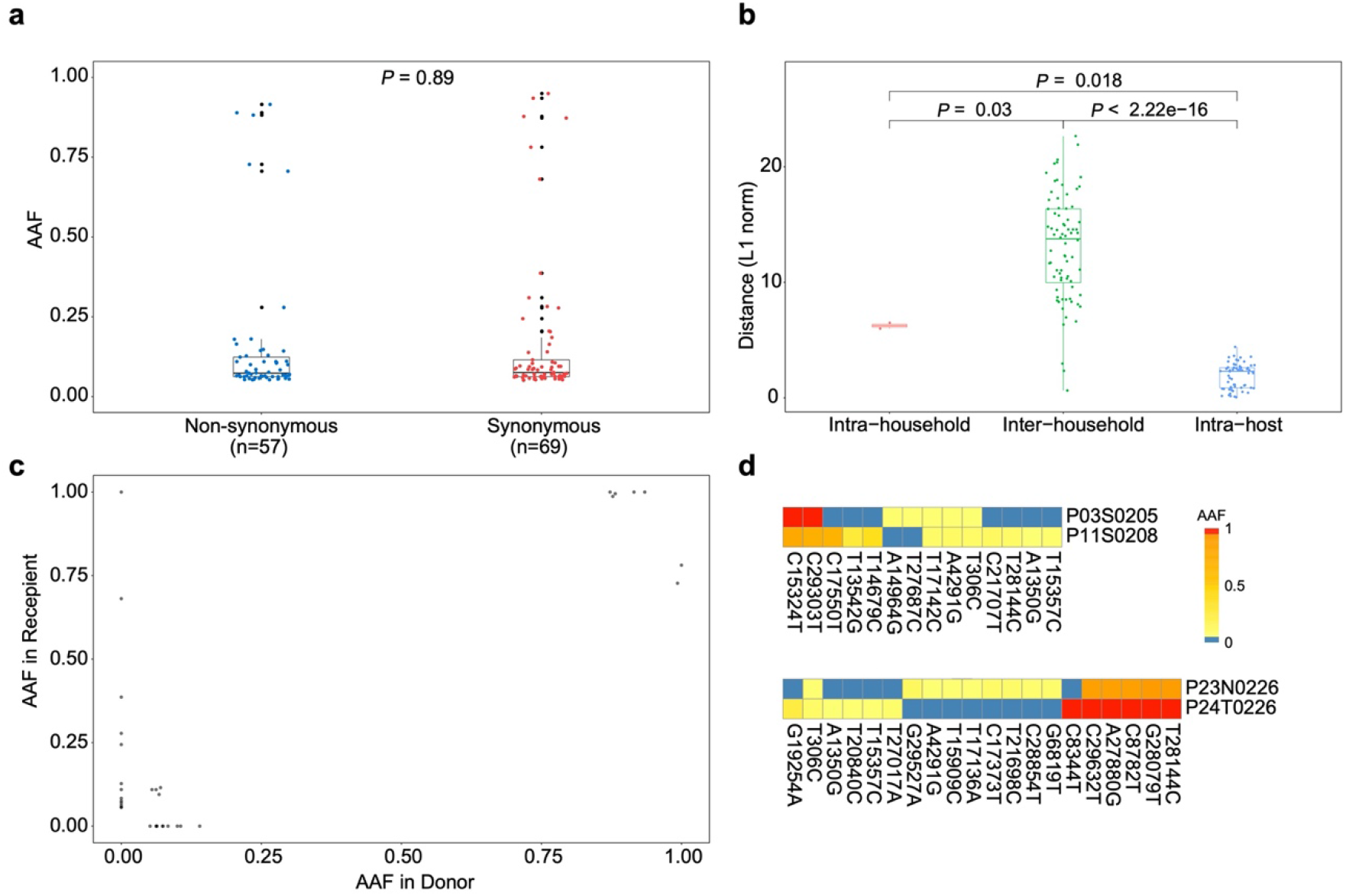
Allele frequency changes of transmission pairs. **a,** Box plots showing the alternative allele frequency (AAF) distribution of synonymous and non-synonymous intra-host variants. **b,** Box plots representing the L1-norm distance distribution among sample pairs. Each dot represents the genetic distance between each sample pair. **c,** The AAF of donor iSNVs in transmission pairs. Allele frequencies under 5% and over 95% were adjusted to 0% and 1, respectively. **d,** Heatmap representing the alternative allele frequencies (AAFs) of consensus and intra-host single nucleotide variants (iSNVs) of the two transmission pairs.

We then estimated the viral genetic distance among samples in a pairwise manner based on their iSNVs and allele frequencies. The samples were firstly categorised into intra-host pairs (serial samples from the same host), intra-household pairs and inter-household pairs (**Figure 2b and Table S4**). As expected, the intra-host pairs had the lowest genetic distance compared to either intrahousehold pairs (Wilcoxon rank sum test, *P* = 0.018) and inter-household pairs (Wilcoxon rank sum test, *P* < 2.22e-16). Interestingly, the genetic distance between intra-household pairs was significantly lower than that of inter-household pairs (**Figure 2b;** Wilcoxon rank sum test, *P* = 0.03), supporting a direct passage of virions among intra-household individuals. Nonetheless, we only observed a few minor variants shared among intra-household pairs, suggesting that the estimated genetic similarity was mostly determined by consensus nucleotide differences (**Figure 2c,d**). Specifically, in one intrahousehold pair (P23 and P24), one patient (P23) contained iSNVs that were coincided with the linked variants, C8782T and T28144C, suggesting that this patient may have been co-infected by genetically distinct viruses. However, the strain carrying C8782T and T28144C was not observed in the intrahousehold counterpart (P24). It is likely that there is a narrow transmission bottleneck allowing only the major strain to be circulated, if P23 was infected by all the observed viral strains before the transmission.

The transmission bottlenecks among intra-household pairs were estimated using a beta binomial model, which was designed to allow some temporal stochastic dynamics of viral population in the recipient [10]. Here, we defined the donor and recipient within the intra-household pairs according to their dates of the first symptom onset. The estimated bottleneck sizes were 6 (P03 and P11) and 8 (P23 and P24) for the two intra-household pairs (**Table S5**). This result is consistent with the patterns observed in many animal viruses and human respiratory viruses [11,12], while the only study reporting a loose bottleneck among human respiratory viral infections [13] was argued as the generic consequence of shared iSNVs caused by read mapping artefacts [14]. The relatively narrow transmission bottleneck sizes is expected to increase the variance of viral variants being circulated between transmission pairs [15]. Even after successful transmission, virions carrying the minor variants are likely to be purged out due to the frequent stochastic dynamics within the respiratory tract [7], which is also consistent with the low diversity and instable iSNV observed among the RT samples.

The observed narrow transmission bottleneck suggests that, in general, only a few virions successfully enter host cells and eventually cause infection. Although the number of transmitted virions is sparse, they can easily replicate in the respiratory tract, given the observed viral replication activities in all the RT sample types and the high host-cell receptor binding affinity of SARS-CoV-2 [16]. The narrow transmission bottleneck also indicate that instant hand hygiene and mask-wearing might be particular effective in blocking the transmission chain of SARS-CoV-2.

In summary, we have characterized and compared SARS-CoV-2 populations of patients within and across households using both metatranscriptomic and hybrid-capture based techniques. Our work (showed an active viral replication activity in the human respiratory tract and the co-existence of genetically distinct viruses within the same host. The inter-host comparison among viral populations further revealed a narrow transmission bottleneck between patients from the same households, suggesting a dominated role of stochastic dynamics in both inter-host and intra-host evolution. The present work enhanced our understanding of SARS-CoV-2 virus transmission and shed light on the (integration of genomic and epidemiological in the control of this virus.

## Materials and methods

### Patient and Ethics statement

Respiratory tract (RT) samples, including nasal swabs, throat swabs, sputum, were collected from 13 COVID-19 patients during the early outbreak of the pandemic (from January 25 to February 10 of 2020). Those patients were hospitalized at the first affiliated hospital of Guangzhou Medical University (10 patients), the fifth affiliated hospital of Sun Yat-sen University (1 patient), Qingyuan People’s Hospital (1 patient) and Yangjiang People’s Hospital (1 patient). The research plan was assessed and approved by the Ethics Committee of each hospital. All the privacy information was anonymized.

### Dataset description

Public consensus sequences were downloaded from GISAID.

### Real-time RT-qPCR and sequencing

RNA was extracted from the clinical RT samples using QIAamp Viral RNA Mini Kit (Qiagen, Hilden, Germany), which was then tested for SARS-CoV-2 using Real-time RT-qPCR. Human DNA was removed using DNase I and RNA concentration was measured using Qubit RNA HS Assay Kit (Thermo Fisher Scientific, Waltham, MA, USA). After human DNA-depletion, the samples were RNA purified and then subjected to double-stranded DNA library construction using the MGIEasy RNA Library preparation reagent set (MGI, Shenzhen, China) following the method used in the previous study [17]. Possible contamination during experimental processing was tracked using human breast cell lines (Michigan Cancer Foundation-7). The constructed libraries were converted to DNA nanoballs (DNBs) and then sequenced on the DNBSEQ-T7 platform (MGI, Shenzhen, China), generating paired-end short reads with 100bp in length. Most samples were also sequenced using hybrid capture-based enrichment approach that was described in previous study [17]. Briefly, the SARS-CoV-2 genomic content was enriched from the double-stranded DNA libraries using the 2019-nCoVirus DNA/RNA Capture Panel (BOKE, Jiangsu, China). The enriched SARS-CoV-2 genomic contents were converted to DNBs and then sequenced on the MGISEQ-2000 platform, generating paired-end short reads with 100bp in length.

### Data filtering

Read data from both metatranscriptomic and hybrid capture based sequencing were filtered following the steps described in the previous research [17]. In brief, short read data were mapped to a database that contains major coronaviridae genomes. Low-quality, adaptor contaminations, duplications, and low-complexity within the mapped reads were removed to generate the high quality coronaviridae-like short read data.

### Profiling of sub-genomic messenger RNA (sgmRNAs)

Coronaviridae-like short reads were mapped to the reference genome (EPI_ISL_402119) using the aligner HISAT2 [18]. Reads spanning the transcription regulatory sequence (TRS) sites of both leader region and the coding genes (S gene, ORF3a, 6, 7a, 8, E, M and N gene) were selected to represent the sgmRNAs. The junction sites were predicted using RegTools junctions extract [19]. The ratio of sgmRNA reads to the viral genomic RNA reads (sgmRNA ratio) was used to estimate the relative transcription activity of SARS-CoV-2.

### Detection of intra-host variants

We defined an intra-host single nucleotide variant (iSNV) as the co-existence of an alternative allele and the reference allele at the same genomic position within the same sample. To identify iSNV sites, paired-end metatranscriptomic coronaviridae-like short read data were mapped to the reference genome (EPI_ISL_402119) using BWA aln (v.0.7.16) with default parameters [20]. The duplicated reads were detected and marked using Picard MarkDuplicates (v. 2.10.10) (http://broadinstitute.github.io/picard). Nucleotide composition of each genomic position was characterized from the read mapping results using pysamstats (v. 1.1.2) (https://github.com/alimanfoo/pysamstats). The variable sites of each sample were identified using the variant caller LoFreq with default filters and the cut-off of 5% minor allele frequency. After filtering the sites with more than one alternative allele, the rest sites were regarded as iSNV sites. All the iSNVs with less than five metatranscriptomic reads were verified using the hybrid capture data (at least two reads). The identified iSNVs were then annotated using the SnpEff (v.2.0.5) with default settings [21].

### Genetic distance

The genetic distance between sample pairs was calculated using L1-norm distance, as defined by the following formula. The L1-norm distance (*D*) between sample pairs is calculated by summing the distance of all the variable loci (*N*). The distance on each variable locus is calculated between vectors (*p* and *q* for each sample) of possible base frequencies (*n* = 4 ¿.

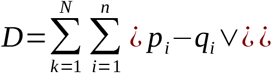

To verify the result, L2-norm distance (Euclidean distance) between sample pairs was calculated. The L2-norm distance *d* (*p,q*) between two samples (*p,q*) is the square root of sum of distance across all the variable loci (*N*), as defined by the following formula.

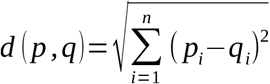

The comparison of genetic distances among sample pair categories was performed using the Wilcoxon rank-sum test.

### Beta binomial model of bottleneck size estimation

A beta-binomial model was used to estimate bottleneck sizes between donors and recipients. Here, the bottleneck size represents the number of virions that pass into the recipient and finally shape the sequenced viral population. The patient with the earlier symptom onset date was defined as the donor, while the other was defined as the recipient. The maximum-likelihood estimates (MLE) of bottleneck sizes were estimated within 95% confidence intervals.

## DATA AVAILABILITY

The data that support the findings of this study have been deposited into CNSA (CNGB Sequence Archive) of CNGBdb with the accession number CNP0001111 (https://db.cngb.org/cnsa/).

## DISCLOSURE STATEMENT

No conflict of interest was reported by the authors

## ACKNOWLEDGEMENTS

This study was approved by the Health Commission of Guangdong Province to use patients’ specimen for this study. This study was funded by grants from The National Key Research and Development Program of China (2018YFC1200100, 2018ZX10301403, 2018YFC1311900), the emergency grants for prevention and control of SARS-CoV-2 of Ministry of Science and Technology (2020YFC0841400), Guangdong province (2020B111107001, 2020B111108001, 2020B111109001, 2018B020207013, 2020B111112003), the Guangdong Province Basic and Applied Basic Research Fund (2020A1515010911), Guangdong Science and Technology Foundation (2019B030316028), Guangdong Provincial Key Laboratory of Genome Read and Write (2017B030301011) and Guangzhou Medical University High-level University Innovation Team Training Program (Guangzhou Medical University released [2017] No.159). This work was supported by the Shenzhen Municipal Government of China Peacock Plan (KQTD2015033017150531). This work was supported by China National GeneBank (CNGB). We thank the patients who took part in this study.

## AUTHOR CONTRIBUTIONS

D.W., Y.X., J.L., W.Z. and J.Z. conceived the study, Y.W., L.Z., and Y.L. collected clinical specimen and executed the experiments. D.W., W.S., X.C. and J.J. analyzed the data. All the authors participated in discussion and result interpretation. D.W., Y.W., and Z.Z. wrote the manuscript. All authors revised and approved the final version.

## SUPPLEMENTARY INFORMATION

**Table S1. Demography and clinical outcomes of COVID-19 patients**

**Table S2. Summary of iSNVs**

**Table S3. Frequency of iSNVs**

**Table S4. Inter-host genetic distance (L1 and L2-norm)**

**Table S5. Bottleneck size of intra-household pairs**

**Figure S1.**
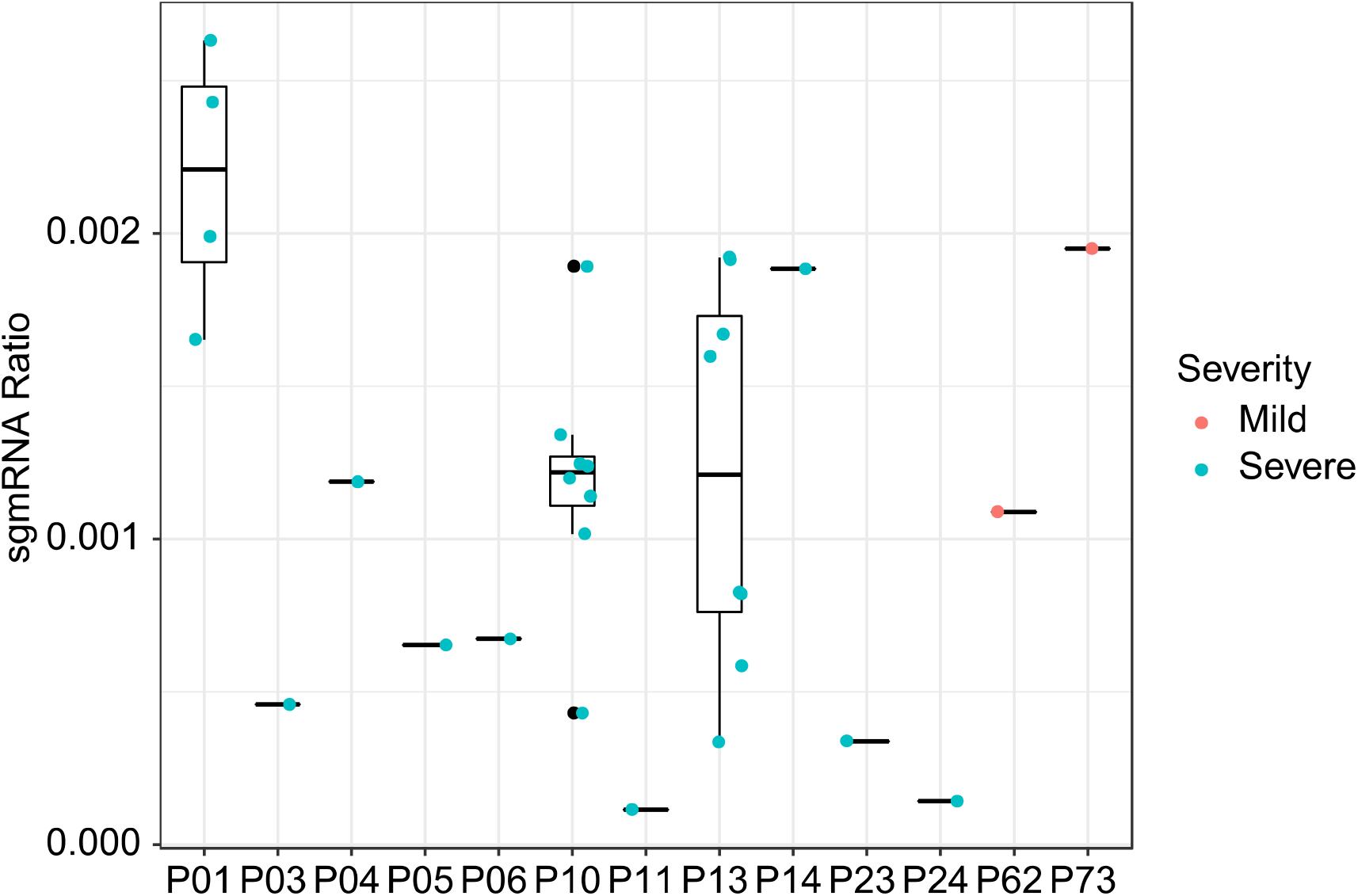
Transcription profile of sub-genomic messenger RNAs (sgmRNAs) of each patient.

**Figure S2.**
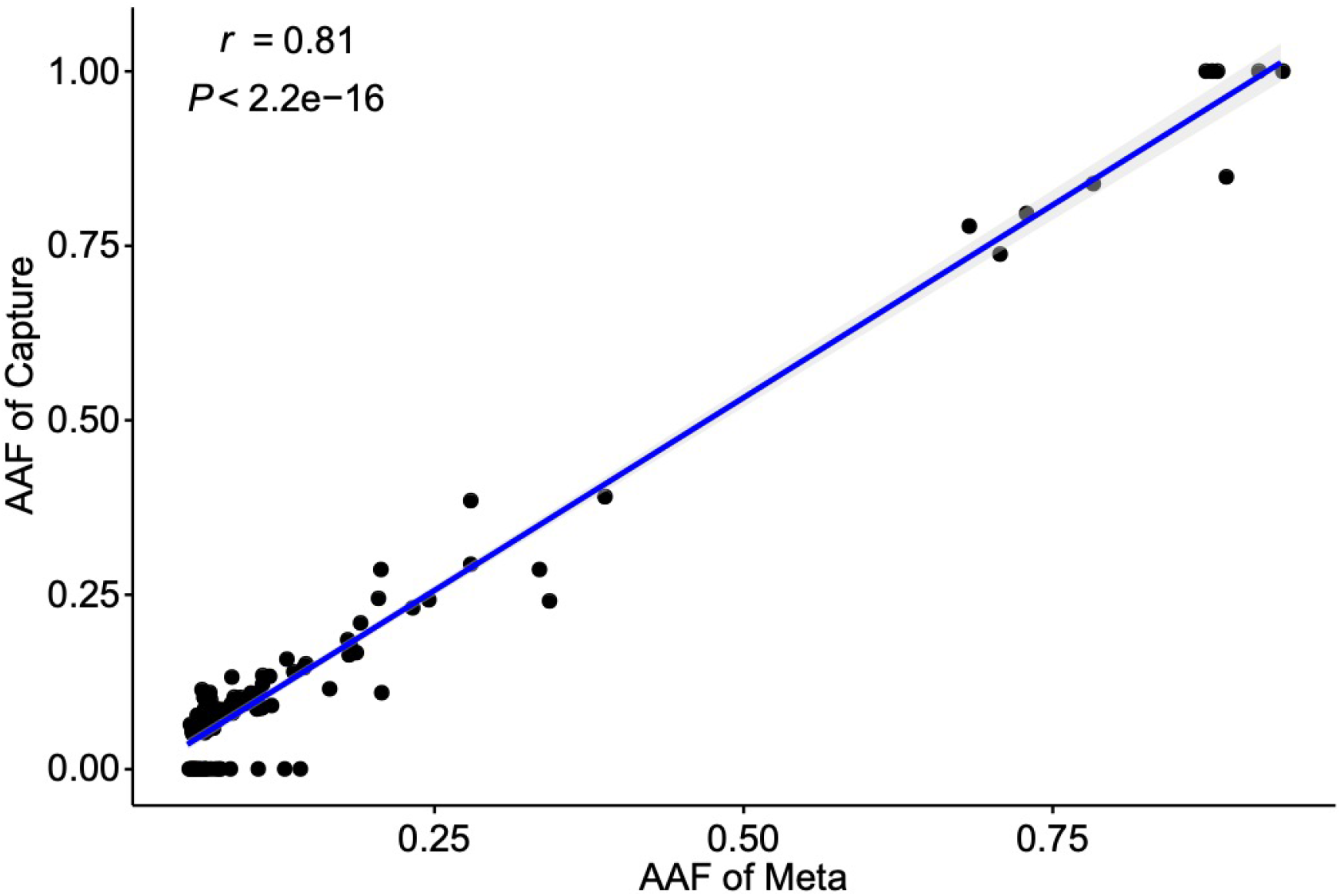
Concordance between minor alternative allele frequencies (AAFs) derived from metagenomic and hybrid capture data.

**Figure S3.**
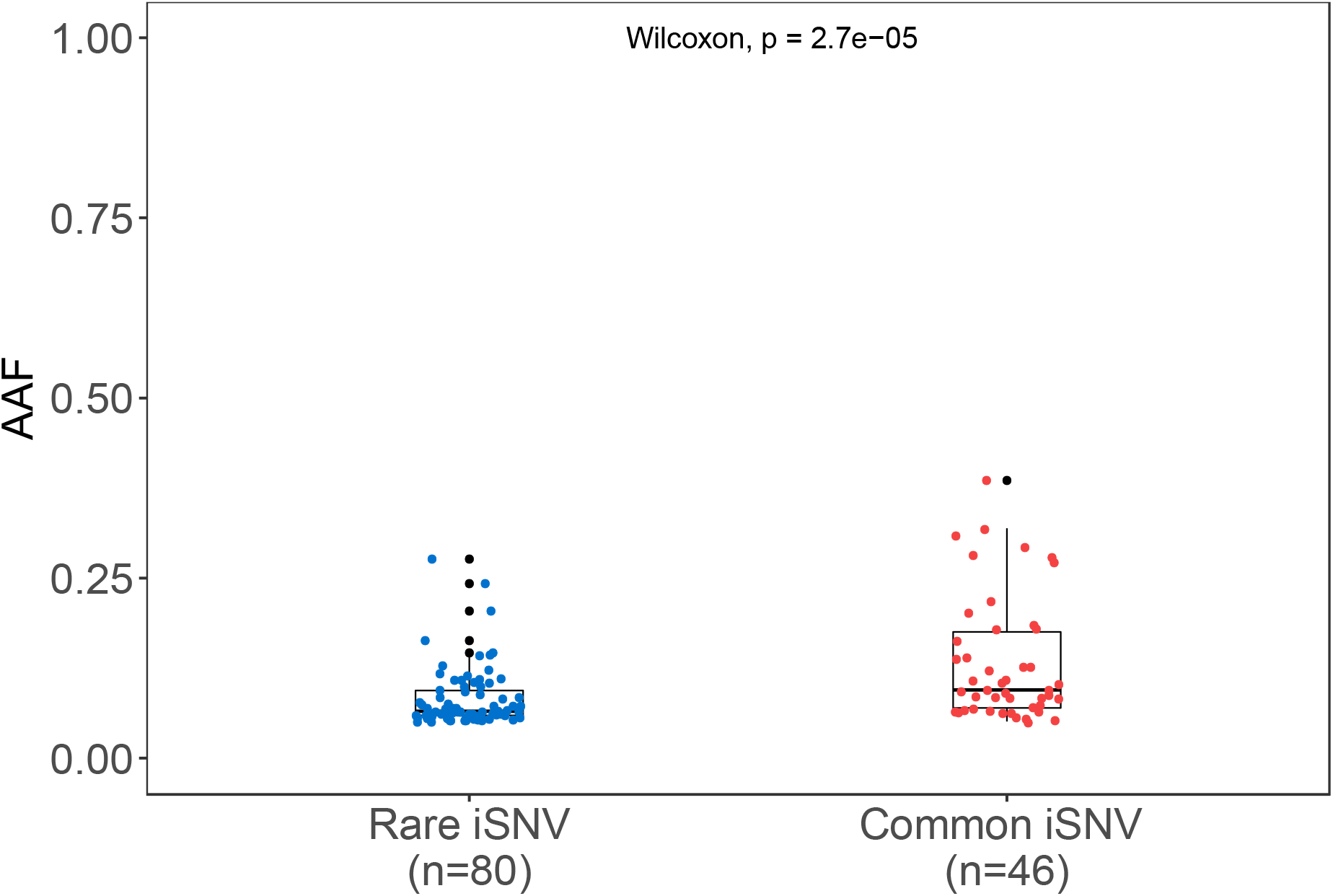
Alternative allele frequency (AAF) distribution of rare and common iSNVs

